# Structural and temporal dynamics analysis on PANoptosis in sepsis: a bibliometric analysis

**DOI:** 10.1101/2025.06.02.657552

**Authors:** Zhihua Li, Di Nie, Lele Yin, Qindan Qin, Chunbo Yang, Rong Li, Xiaoming Gao, Xiangyou Yu, Yi Wang

## Abstract

PANoptosis, as a new type of programmed cell death, is characterized by pyroptosis, apoptosis and necroptosis, and is a key mechanism causing a variety of inflammatory diseases. Despite the growing number of studies indicating the crucial role of PANoptosis in sepsis, there has been no bibliometric analysis of the research hotspots and trends in this field. Therefore, this study aims to explore the history, research hotspots and emerging trends of PANoptosis in sepsis related research in the past 20 years from the perspective of structure and temporal dynamics. The articles related to PANoptosis in sepsis were retrieved from the Web of Science Core Collection (WoSCC) database from 2000 to 2024. CiteSpace and HistCite were used to analyse the historical features, the evolution of active topics, and emerging trends about PANoptosis in sepsis. 6165 original articles and reviews on PANoptosis in sepsis were included in the bibliometric analysis. In the last 20 years, the number of published documents is increasing year by year and reaches a peak in 2022. At the same time, many activation themes have emerged at different times, as evidenced by a total of 96 categories, 865 keywords and 629 reference bursts. Keyword clustering anchored eight emerging research subfields, namely 0# oxidative stress, 1#pyroptosis, 2#sepsis-associated encephalopathy, 3#acute kidney injury, 4#immunosuppression, 5#necroptosis, 7#lung injury and 8#extracellular vesicles. The keyword alluvial map shows that the most persistent research concepts in this field are phosphatase. And the emerging keywords are toll_like_receptors, regulatory T cells, recognition, etc. In a timeline visualization based on the time span of the citations, we find five relevant emerging topics, namely 1# immunosuppression, 2# sepsis-induced cardiomyopathy, 3# pyroptosis, 4# acute kidney injury, and 13# COVID-19. Our study provides a comprehensive bibliometric analysis and summary about the current status and trends on PANoptosis in sepsis, which will aid researchers in conducting further scientific research in this field.

## 1. Introduction

Sepsis is defined as a life-threatening organ dysfunction triggered by the host’s dysfunctional systemic inflammatory response and immune response to infection^1^. According to a global epidemiological survey, there were 48.9 million cases of sepsis in the world, including 11 million cases of sepsis related deaths, accounting for 19.7% of the total number of global deaths^2^. It is estimated that the annual medical cost of treating 230,000 sepsis patients in the ICU in China is about 4.6 billion US dollars, and this high medical expenditure puts a heavy economic burden on society and families^3^.

During the occurrence and development of sepsis, different forms of cell death, such as apoptosis, necroptosis, and pyroptosis^4–6^, are regulated to defend against pathogen invasion and maintain homeostasis. Previous studies believed that various forms of cell death exist independently, but more and more studies have shown that there is a wide range of crosstalk between various forms of cell death effective molecules^7–10^, which promotes the establishment of a new form of cell death, namely PANoptosis^11^. It is triggered by specific stimuli and regulated through the PANoptosome complex^12^. Furthermore, several studies have indicated the pivotal role of PANoptosis in multiorgan dysfunction resulting from sepsis. Inhibiting the degree of PANoptosis can improve the lung injury^13^ and myocardial injury^14^ caused by sepsis. Although some progress has been made in the study of PANoptosis in sepsis, there are still some problems to be explored. Therefore, we intuitively analyzed the research hotspots of PANoptosis and sepsis in the past 20 years through the bibliometrics method in WoSCC database.

In contrast to conventional reviews grounded in academic perspectives, bibliometric reviews, which rely on qualitative transformations in academic outputs, offer a more impartial and exhaustive portrayal of the historical trajectory, focal research areas, and evolutionary patterns within a given discipline^15–17^. Various bibliometric instruments have been deployed within the realm of scientometrics, such as CiteSpace^18^, CitNetExplorer^19^, VOSviewer^20^, and HistCite^21^, to analyze and delineate the profile of an academic domain.

In this study, the software applications CiteSpace (version 5.8 R3), HistCite Pro 2.1, and the alluvial diagram generator were utilized to conduct an evaluation of the bibliographic records pertaining to sepsis-induced PANoptosis research. The content of this article encompasses, firstly, summarizing the state of research on PANoptosis in sepsis. Secondly, highlighting the significant contributions of key literature within this field. Thirdly, summarizing the prevalent topics of research in this area, and finally, revealing emerging trends for future research endeavors.

## 2. Methods

### 2.1 Data collection and statistics

The Thomson Reuters Web of Science Core Collection (WoSCC) encompasses over 12,000 prestigious academic journals that are universally acknowledged as significant within the international scholarly community. This research utilizes the WoSCC as the object database. The search criteria employed were: (((((((((((((((((((TS = (Sepsis)) OR TS = (“Bloodstream Infection”)) OR TS = (“Bloodstream Infections”)) OR TS = (“Infection, Bloodstream”)) OR TS = (Septicemia)) OR TS = (Septicemias)) OR TS = (“Blood Poisoning”)) OR TS = (“Blood Poisonings”)) OR TS = (“Poisonings, Blood”)) OR TS = (“Poisoning, Blood”)) OR TS = (“Severe Sepsis”)) OR TS = (“Sepsis, Severe”)) OR TS = (Pyemia)) OR TS = (Pyemias)) OR TS = (Pyaemia)) OR TS = (Pyaemias)) OR TS = (Pyohemia)) OR TS = (Pyohemias)) AND ((((((((((((((TS = (PANoptosis)) OR TS = (Pyroptosis)) OR TS = (Pyroptoses)) OR TS = (“Pyroptotic Cell Deaths”)) OR TS = (“Caspase-1 Dependent Cell Death”)) OR TS = (“Caspase 1 Dependent Cell Death”)) OR TS = (Apoptosis)) OR TS = (“Classical Apoptosis”)) OR TS = (“Classic Apoptosis”)) OR TS = (“Classic Apoptoses”)) OR TS = (“Programmed Cell Death, Type I”)) OR TS = (“Cell Death, Programmed”)) OR TS = (“Programmed Cell Death”)) OR TS = (“Caspase Dependent Apoptosis”)) OR TS = (Necroptosis)). The search was conducted over the period spanning from 2000 to 2024. The retrieved literature records were downloaded and archived in plain text format, specifically as “Full Record and Cited References,” serving as a representative sample for the data analyzed in this study. Ultimately, a dataset comprising 6,165 literature entries was compiled and designated as DATA. Additionally, original data pertaining to the publication’s country/region, institution, journal, author, article type, and other pertinent details were gathered and analyzed using EXCEL (WPS 21019) for statistical purposes.

### 2.2 Tools for bibliometric analysis

#### 2.2.1 CiteSpace

##### The co-occurrence networks

Scientific research inherently demands extensive collaboration, and an in-depth analysis of scientific cooperation can provide profound insights into the status and dynamics of research within a specific scientific discipline. This analysis can be encapsulated within three fundamental dimensions: authorship, institutional affiliation, and nationality. The collaborative endeavors and scientific constructs associated with these dimensions can be visualized as a co-occurrence network when datasets comprising articles from a particular research domain are imported into the CiteSpace software. CiteSpace utilizes color-coded nodes and edges to differentiate various components within the merged network, assigning annual colors based on the chronological timeline of the dataset. The color located at the periphery of the network signifies the initial year in which a co-occurrence link was established. The nodes are composed of multi-colored “tree rings,” where the thickness of each ring represents the frequency of co-occurrence in a specific year. Notably, a red ring signifies a citation burst in a particular year, which is indicative of a substantial and abrupt increase in citations during that specific period. Conversely, a purple ring is employed to delineate the level of node centrality within the network, with a node exhibiting high betweenness centrality being particularly significant as it functions as a crucial bridge connecting other nodes within the network.

##### Burst detection

According to Jon Kleinberg, a sequence of documents, such as emails or articles, possesses a distinct and coherent topic for a finite duration, after which it undergoes a phase of decline. By employing specific text data mining algorithms, this temporal evolution of topics can be identified and represented through the phenomenon of “bursts of activity.” Building upon Kleinberg’s foundational algorithm, Chen et al. defined citation bursts as an indicator of topics that are currently the subject of active discussion within a research community. Citation bursts refer to the detection of emergent events that may persist for several years or even a solitary year, marking periods of intense engagement with particular topics. CiteSpace provides functionality for detecting citation bursts across various disciplines, keywords, and references. The occurrence of citation bursts serves as empirical evidence that a particular discipline, keyword, or reference is associated with a notable surge in citations, thereby indicating that such topics, keywords, or references are garnering substantial attention and scrutiny from the broader scientific community.

##### The cluster analysis

CiteSpace provides three clustering algorithms that harness titles, abstracts, and keywords to organize publications into conceptual clusters, each exhibiting unique research characteristics. The resultant cluster map, influenced by the predefined slicing parameters, visually represents the evolution of these conceptual clusters across various time periods. Additionally, timeline mapping offers a precise depiction of the emergence and decline of individual clusters, as well as the interconnected nodes that link these clusters to others within the broader research landscape.

The detailed methodology employed for this analysis is as follows: Initially, data pertaining to cell death in sepsis was imported into the CiteSpace software, specifically version 6.2.R4. The “Time slice” parameter was configured to encompass the period from 2000 to 2024, with each slice representing one year. For the purposes of analysis, “Author keyword (DE R3)” was selected. Subsequently, the “Time slice” was adjusted to span from 2004 to 2024, maintaining the same slicing interval. For data extraction, we selected the etymological sources comprising “Title,” “Abstract,” “Author keyword (DE),” and “Keyword +.” The node types were specified as required, while all other settings were retained at their default values. Subsequently, the software automatically generated knowledge graphs that illustrated collaboration networks among countries (regions), institutions, and authors. These graphs were then manually refined to enhance clarity and aesthetic quality. Employing an identical methodology, keyword cluster graphs were created, with “keyword” designated as the node type. Distinct time slices were defined for this purpose: 2000- 2006, 2007-2012, 2013-2018, and 2019-2024. When generating cluster maps for references, the Layout TAB within the Control Panel was accessed, and the Timeline TAB was selected to produce a citation timeline map in Timeline View. Furthermore, the Burst TAB in the Control Panel was utilized to execute the View command, thereby generating burst maps that focused specifically on keywords, categories, or references.

#### 2.2.2 HisCite

The HistCite Pro 2.1 software possesses the capability to visualize digital citation networks and identify the most influential documents, thereby empowering researchers to expeditiously ascertain the publications that are most frequently cited within a given field. This software assigns scores to articles based on two metrics: the Local Citation Total Score (LCS) and the Global Citation Total Score (GCS). Specifically, LCS reflects the citation frequency within the software’s dataset, whereas GCS represents the citation frequency within the expansive Web of Science Core Collection (WoS CC) database. In this study, a dataset comprising 6,165 research articles focused on cell death in sepsis, designated as DATA, was imported into HistCite Pro 2.1. With the “Limit” parameter set to 30 and all other settings retained at their default values, the “Make a chart” option was selected to create a citation map of the research field pertaining to cell death in sepsis. This citation map facilitated the swift identification of seminal literature within the domain.

#### 2.2.3 The alluvial generator

Alluvial flow maps are specifically crafted to elucidate temporal patterns within dynamically evolving networks. To produce such a map, we employed the CiteSpace software to generate a series of individual networks that represented the co-occurrence of keywords. These networks were subsequently exported to the debris flow generator, accessible via the URL http://www.mapequation.org/apps/AlluvialGenerator.html. In this methodology, each keyword was treated as a node, and nodes were clustered within each temporal slice, with each cluster constituting a module. Across various temporal slices, nodes underwent processes of splitting and merging to form novel modules, with the most recent module emerging from the intersection of previously existing nodes.

## 3. Results

### 3.1. The historical features of the literature on Cell death in sepsis

#### 3.1.1. Distribution of publications

The quantitative variation of scientific literature at a particular temporal node can offer insights into the accumulation of knowledge within a specific research domain and provide crucial quantitative parameters for understanding the progression of this field. A comprehensive search yielded a total of 2601 articles pertinent to cell death in sepsis, encompassing 55,478 research articles, 687 review papers, contributions from 23,484 authors, involvement of 4051 institutions, and publication across 1212 journals within 101 scientific disciplines (Table 1).

**Table 1.** The distribution of publications.

| Categories | Publication | Articles | Review | Authors | Institutions | Journals | Subject categories |
| --- | --- | --- | --- | --- | --- | --- | --- |
| Amount | 6165 | 5478 | 687 | 23484 | 4051 | 1212 | 101 |

The results of the annual study are shown in Figure 1A. In 2000, a total of 52 papers related to cell death in sepsis were published. Sixty papers were published in 2001. From 2000 to 2003, the number of papers was low, from 2004 to 2017, the number of papers increased rapidly, and the number of papers increased further after 2018, and reached a peak in 2022.

**Figure 1.**
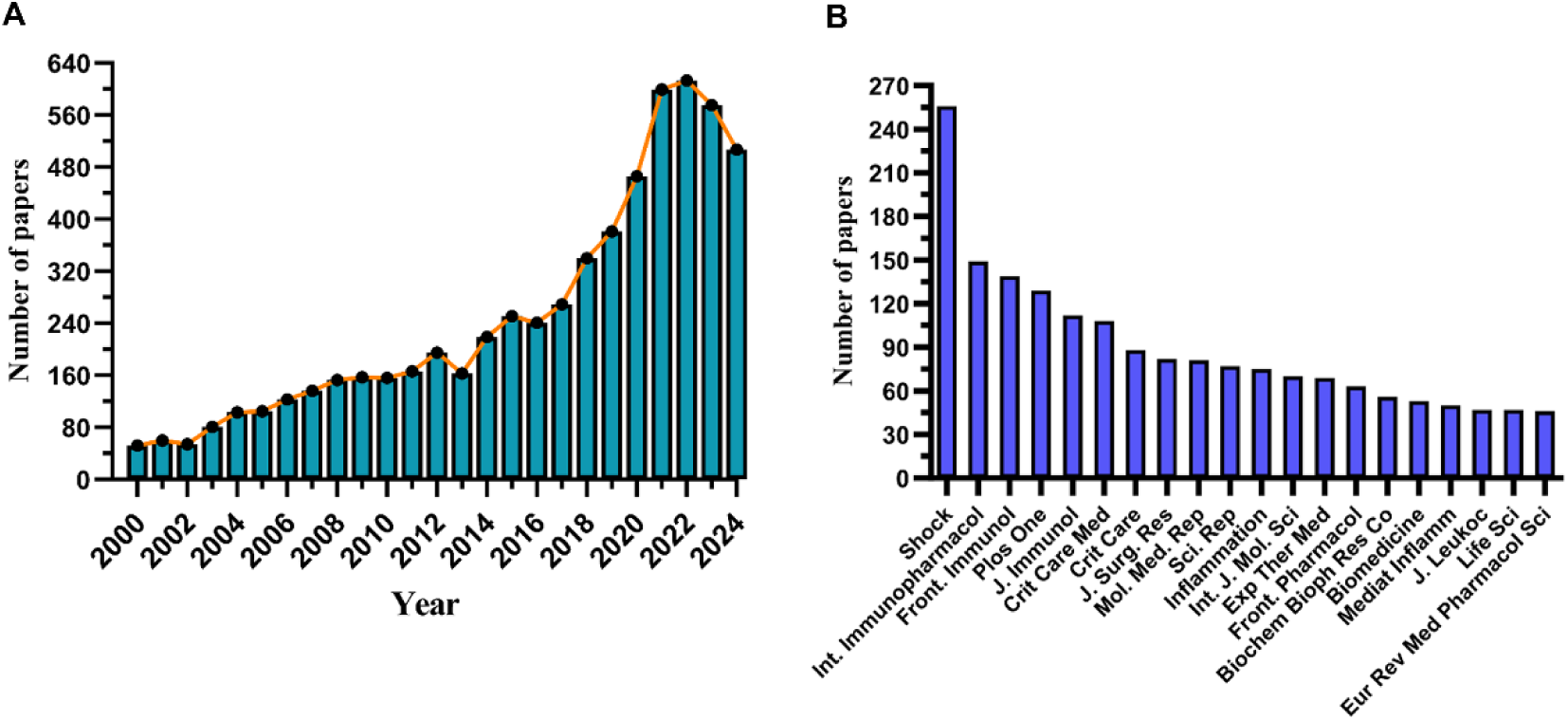
A: Annual number of publications (number of publications per year). B: The top 20 fruitful journals.

Shock ranked first in publication volume (256), followed by international immunopharmacology (149) and frontiers in immunology (139). The top 20 journals with the most results are listed in Figure 1B, which researchers can refer to when considering submissions. These findings indicate that PANoptosis have gained more attention from scholars in recent years and are becoming a significant focus of sepsis.

#### 3.1.2. The vein of research on Cell death in sepsis

The citation cocitation chart delineates the literature relationships within the field of cell death in sepsis over the past two decades (Figure 2). This network comprises a total of 1828 nodes and 8787 links, indicating extensive interconnections among the publications in this research domain. Analogous to a vast tree, the earlier literature spanning from 2000 to 2010 forms the root system of the field, with grey-marked nodes exhibiting dense connectivity and rich inter-node links, serving as the foundational nutrients for the sustained development of the field. During the middle period (2011-2017), the blue-marked nodes gradually diverge to establish the principal branches of the study. In the subsequent period (2018-2024), these nodes further branch out and cluster more tightly, signaling the concentration and differentiation trends within the research field. In these literatures, Singer M (2016)^22^, Rudd KE (2020)^23^, Cecconi M (2018)^24^, Peerapornratana S (2019)^25^, Poston JT (2019)^26^, Li N (2019)^27^, Fleischmann C (2016)^28^, Hotchkiss RS (2003)^29^, Rhodes A (2017)^30^, van der Poll T (2017)^31^, These 10 papers were cited 422, 261, 159, 143, 115, 114, 105, 101, 101 and 96, respectively, and occupy an important position in the field of cell death in sepsis research. The concentration and differentiation of this research cluster will be more clearly displayed in the subsequent reference time graph. In addition, we mapped the citation history of research articles using HisCite Pro 2.1. These landmark papers are highlighted in Table 2, where the top three papers are Medical progress: The pathophysiology and treatment of sepsis., Sepsis-induced apoptosis causes progressive profound depletion of B and CD4+ T lymphocytes in humans and apoptosis and caspases regulate death and inflammation in sepsis. The larger the node, the more important the reference, the more node connections, and the higher the intermediate centrality of the node. Using these two methods, we not only visually see the citation context structure of the literature, but also focus on the high-contribution literature in the field.

**Figure 2.**
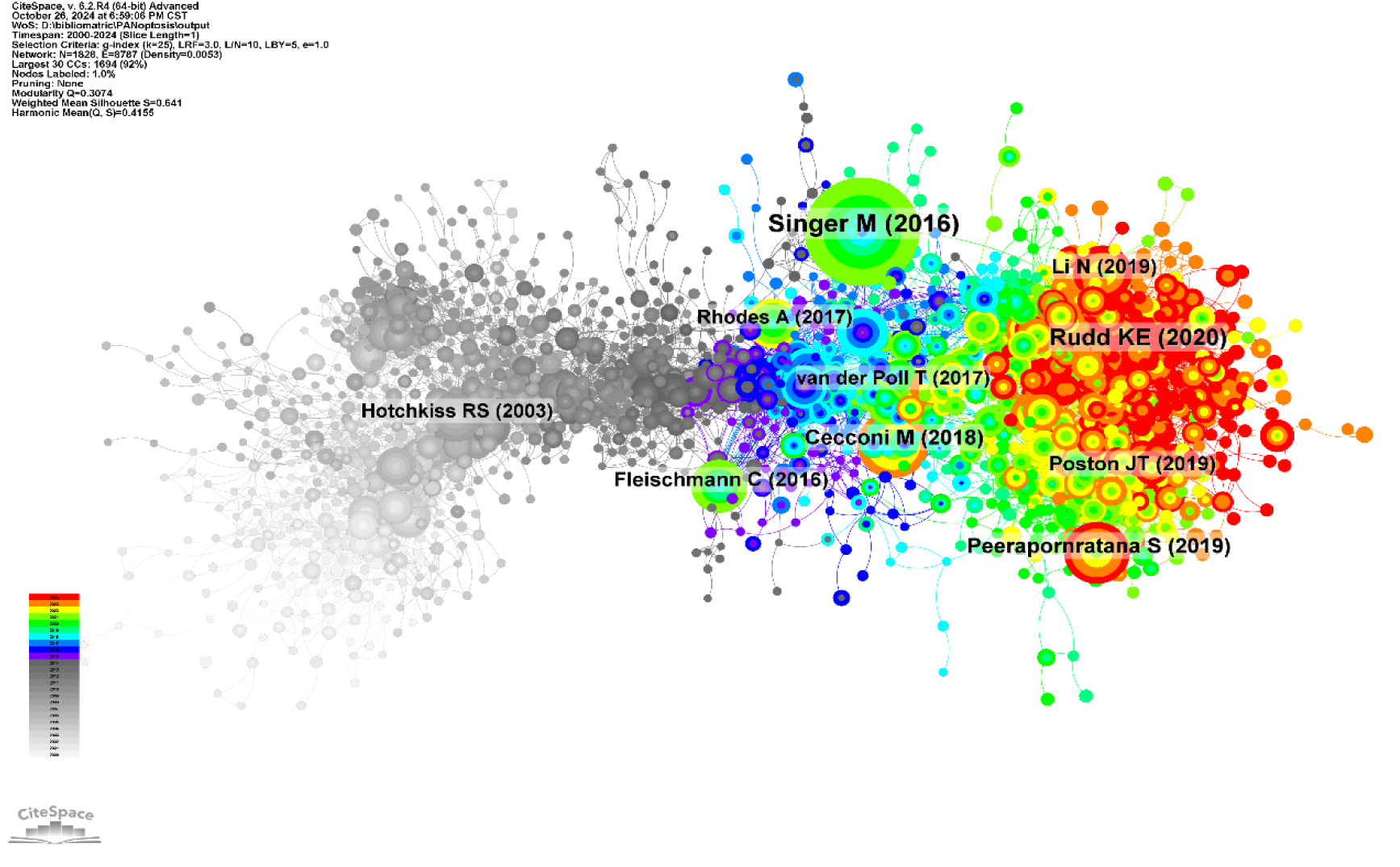
The citation co-occurrence network.

**Table 2.**
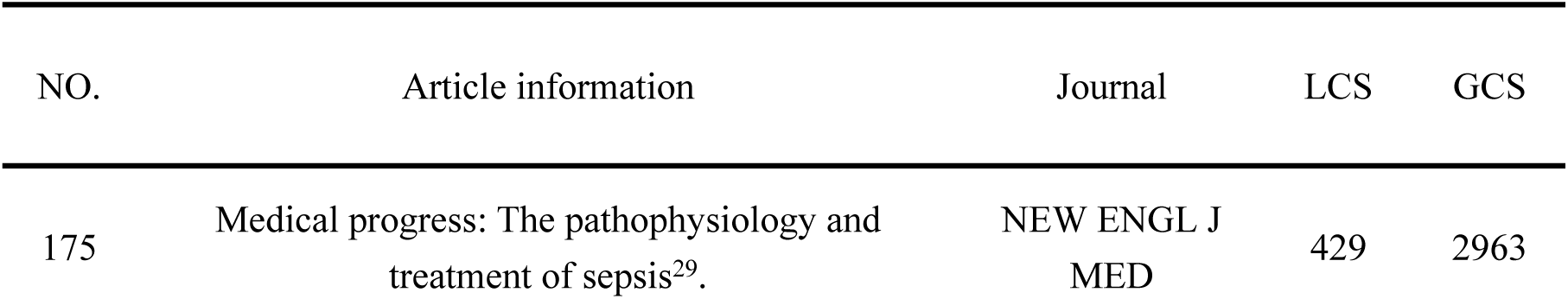
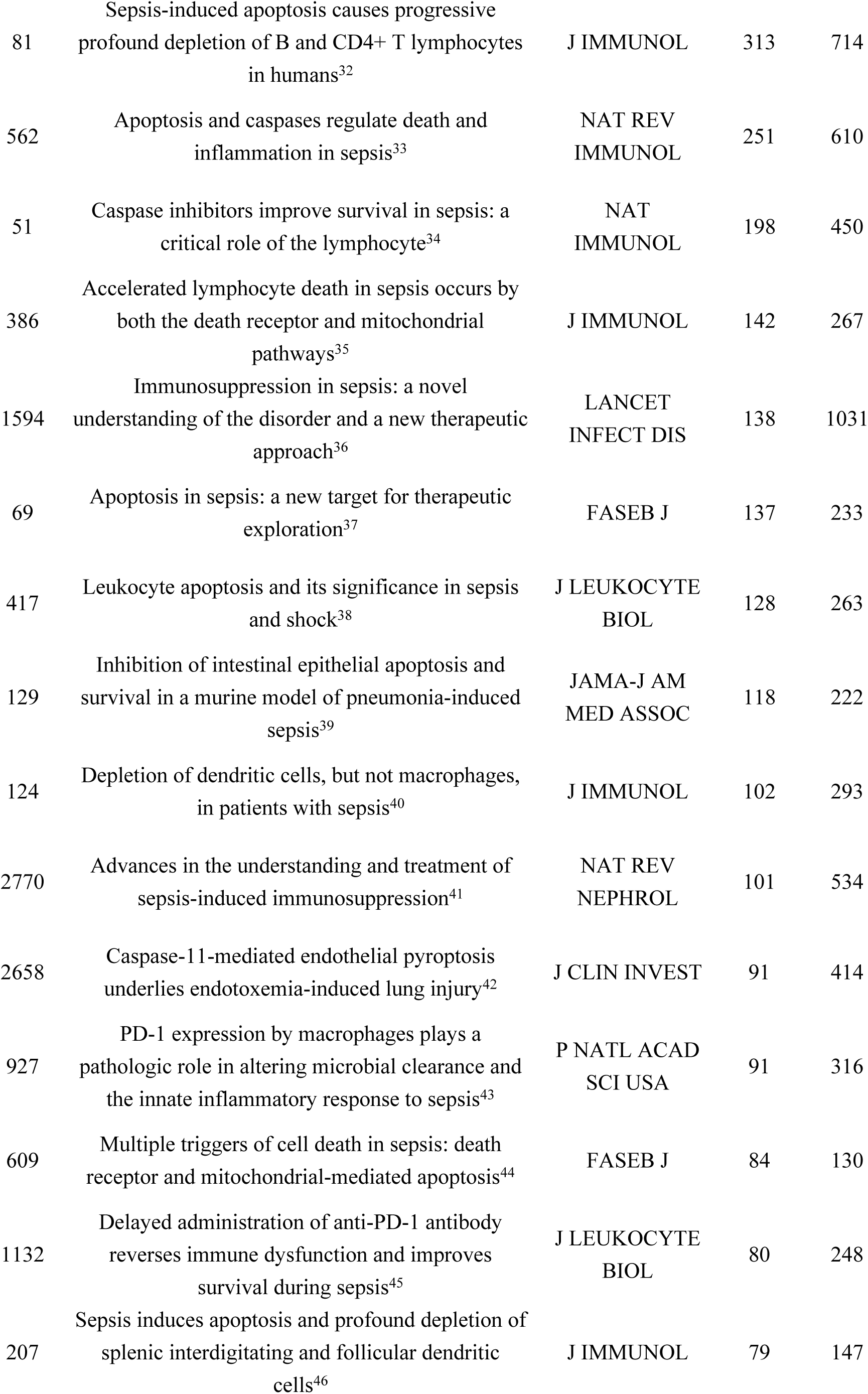
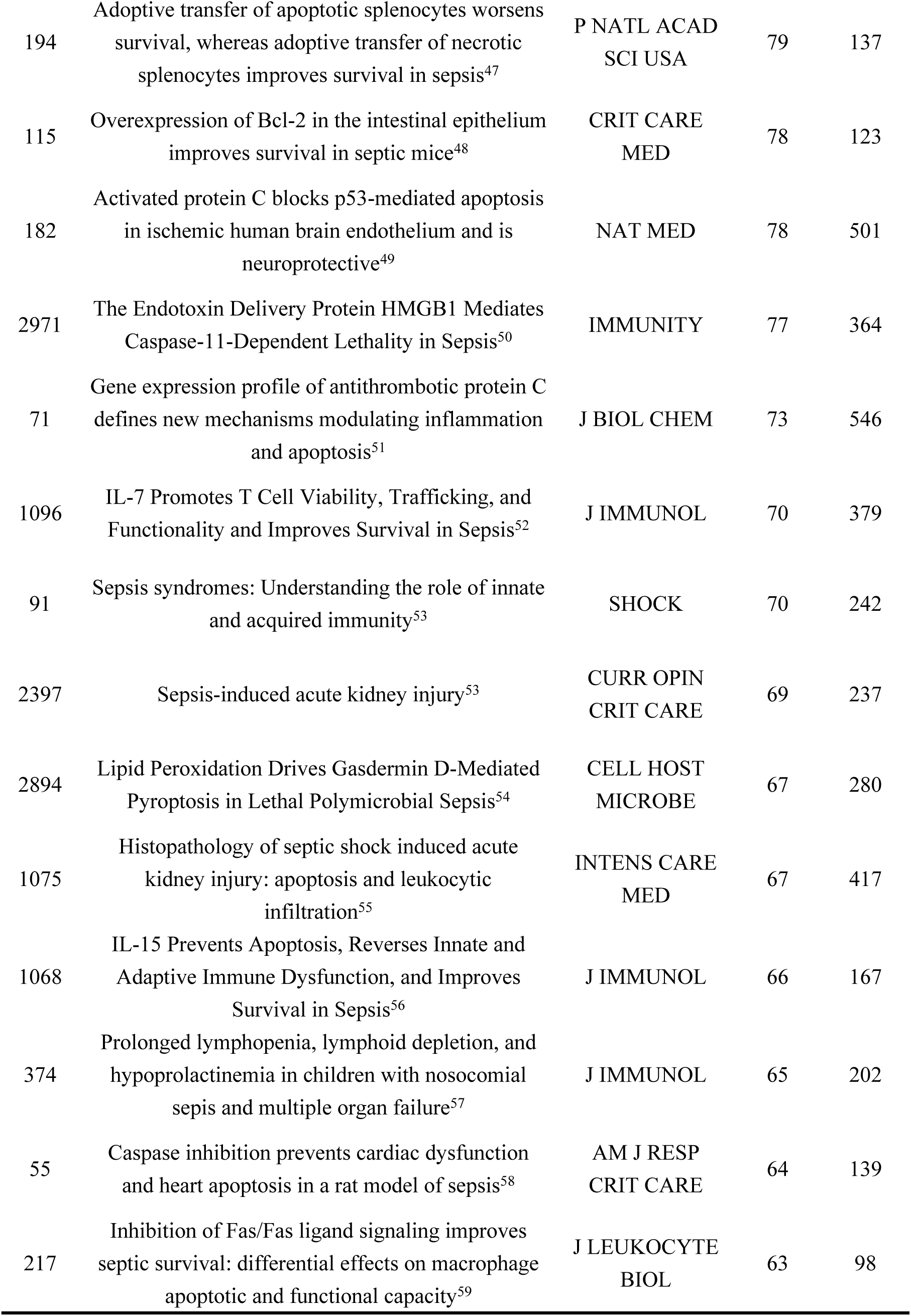
The information of the top 30 literature sorted by LCS score.

#### 3.1.3. Scientific cooperation

As illustrated in Figure 3 and Table S1, the substantial number of nodes and abundant links demonstrate robust scientific collaboration across three dimensions: country, institution, and author. The National Cooperative Network comprises 86 nodes and 481 connections, encompassing nodes from China, the United States, Germany, Japan, and South Korea (Figure 3A). The Institutional Cooperative Network features 599 nodes and 838 connections, with nodes primarily representing Central South University, Shanghai Jiao Tong University, Wuhan University, and Southern Medical University (Figure 3B). The Author Collaboration Diagram is presented in Figure 3C, revealing that Wang Ping, Coopersmith Craig M., Hotchkiss Richard S., Ayala Alfred, and Chung Chun-Shiang have published the highest number of papers in this field. The dense interconnections reflect extensive scientific collaboration among researchers. Notably, a clustering effect is observed among the nodes representing Wang Ping and Aziz Monowar, forming one cluster, while nodes such as Coopersmith Craig M. and Liang Zhe constitute another cluster.

**Figure 3.**
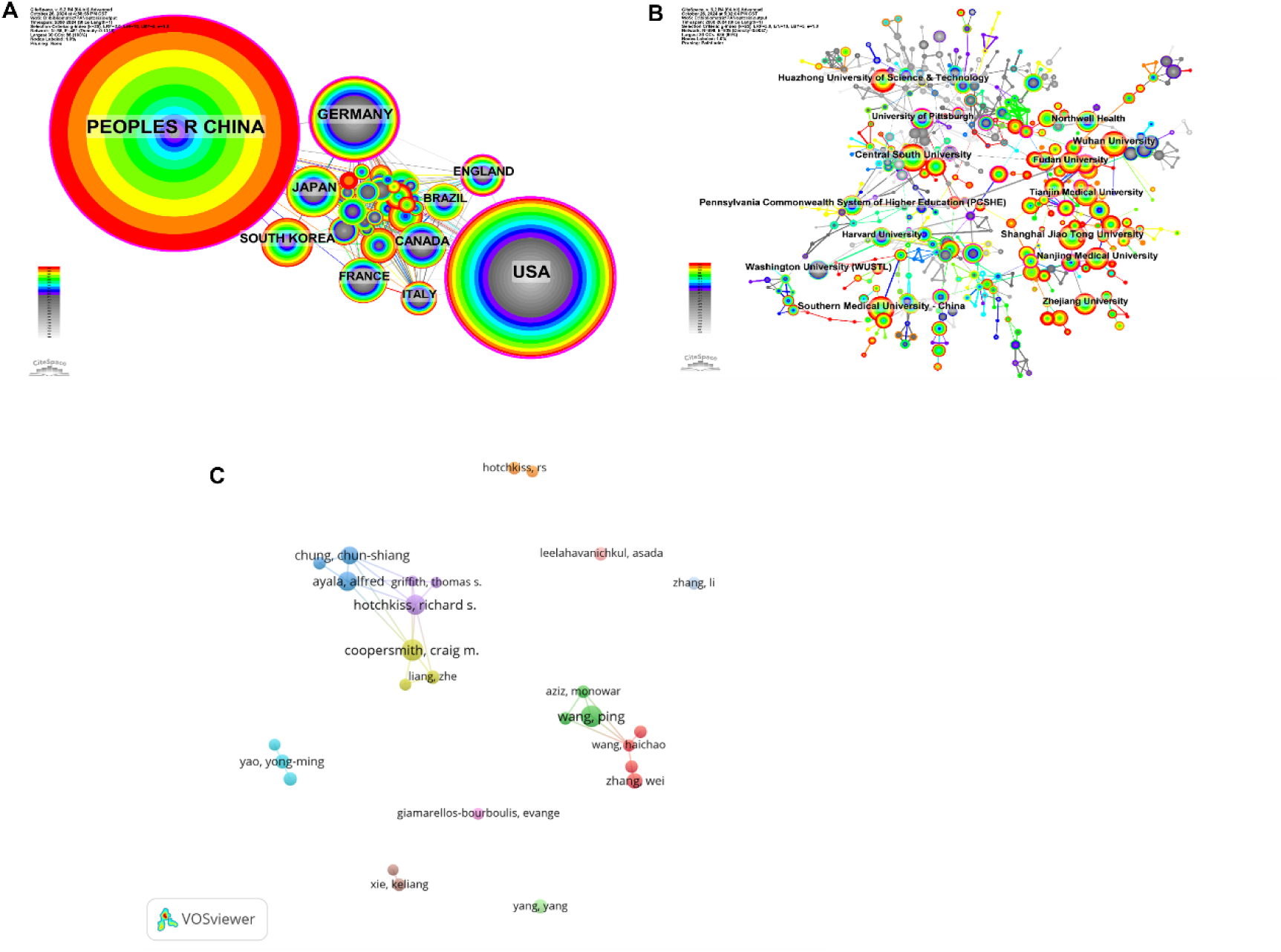
The scientific cooperation network. A: Country cooperation. B: Institution cooperation. C: Author cooperation.

### 3.2. Variation of the most active topics

#### 3.2.1. Subject category burst

From 2000 to 2024, a total of 96 citations underwent significant bursts across 101 pertinent disciplines. The blue line demarcates this temporal interval, while the red line segment denotes the span of the surge in subject categories, with the corresponding years of commencement and termination indicated. Figure 4 depicts the top 50 categories exhibiting high burst intensities at various points in time. CRITICAL CARE MEDICINE experienced the peak burst period between 2000 and 2011, with a burst intensity of 69.88. Notably, there has been a diversification of subject categories over time, including GASTROENTEROLOGY & HEPATOLOGY (2001-2013), PATHOLOGY (2009-2014), UROLOGY & NEPHROLOGY (2014-2016), PSYCHIATRY (2015- 2019), BIOTECHNOLOGY & APPLIED MICROBIOLOGY (2020-2022), and NANOSCIENCE & NANOTECHNOLOGY (2022-2024). These abrupt shifts in disciplinary focus throughout the timeline underscore the multidisciplinary nature of the field. Furthermore, 20 emerging disciplines have been identified from 2024 onwards (Table S2), with the top three being PHARMACOLOGY & PHARMACY (2023-2024), CHEMISTRY, MULTIDISCIPLINARY (2021-2024), and FOOD SCIENCE & TECHNOLOGY (2021-2024).

**Figure 4.**
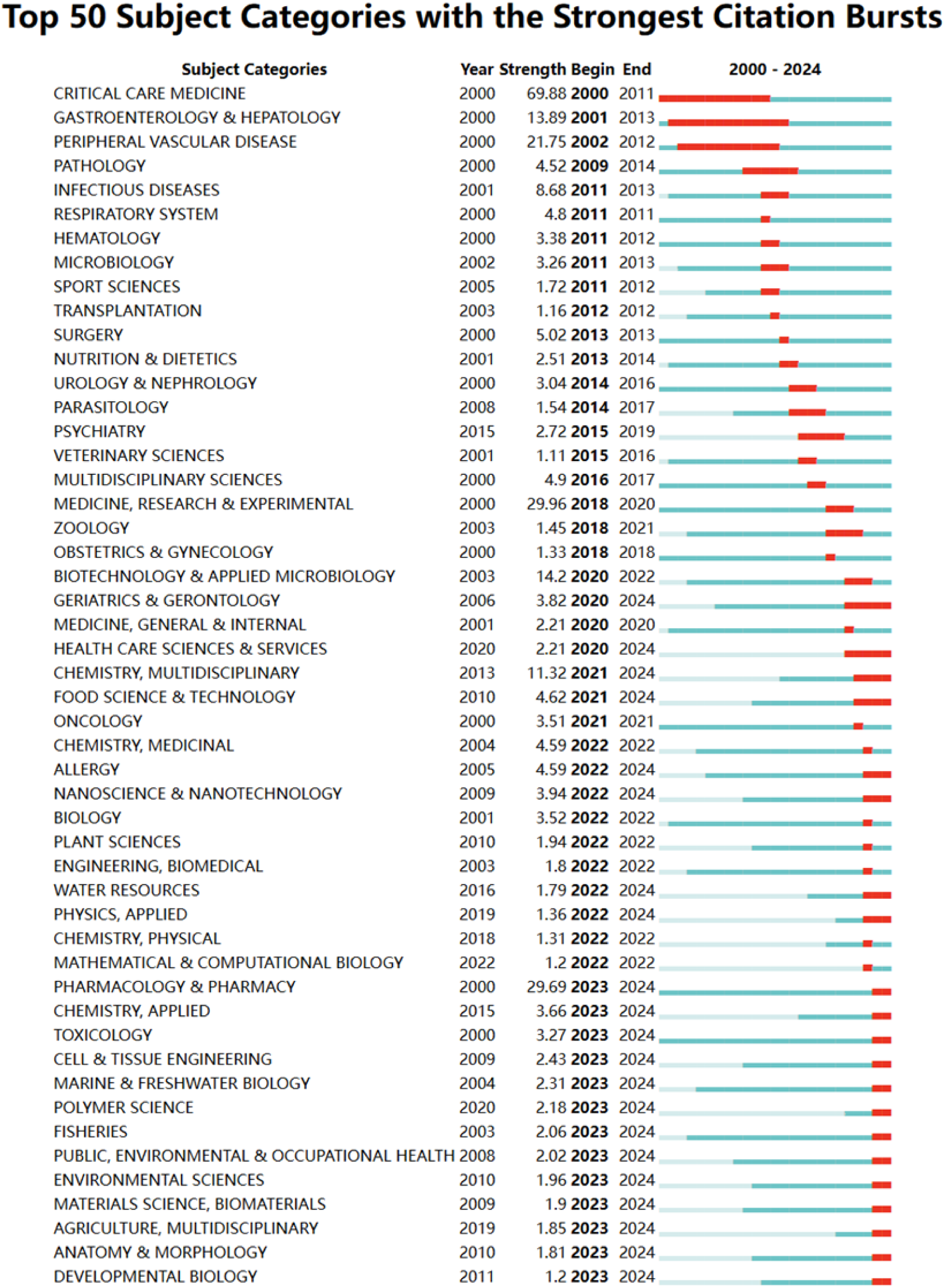
The top 50 categories with high outbreak intensity at different times.

#### 3.2.2. Keywords burst

Keyword burst detection is a method used to identify high-frequency keywords within a specific time frame, aiding researchers in analyzing the evolution of PANoptosis-related cell death. At a finer level of granularity, we investigated the outbreak patterns of keywords to uncover the dynamic content of cell death research in the field of sepsis across the entire study period (2000-2024). A total of 865 keywords exhibited bursts at various time points, with the top 50 keywords demonstrating the highest burst intensities presented in Figure 5. Notably, “tumor necrosis factor” 5exhibited the highest burst intensity (58.24) from 2000 to 2011, followed by “nitric oxide synthase” with a burst intensity of 29.49 between 2001 and 2012, and “endotoxin” with a burst intensity of 27.62 from 2000 to 2012. Furthermore, we focused on the 20 keywords that continued to burst in 2024, as they may represent potential future research hotspots in this field. Specifically, the NLRP3 inflammasome demonstrated a burst intensity of 25.51 between 2021 and 2024, gasdermin D showed a burst intensity of 22.29 within the same period, myocardial injury exhibited a burst intensity of 18.33 from 2020 to 2024, and autophagy had a burst intensity of 16.08 between 2019 and 2024 (see Supplemental Material Table S2 for details).

**Figure 5.**
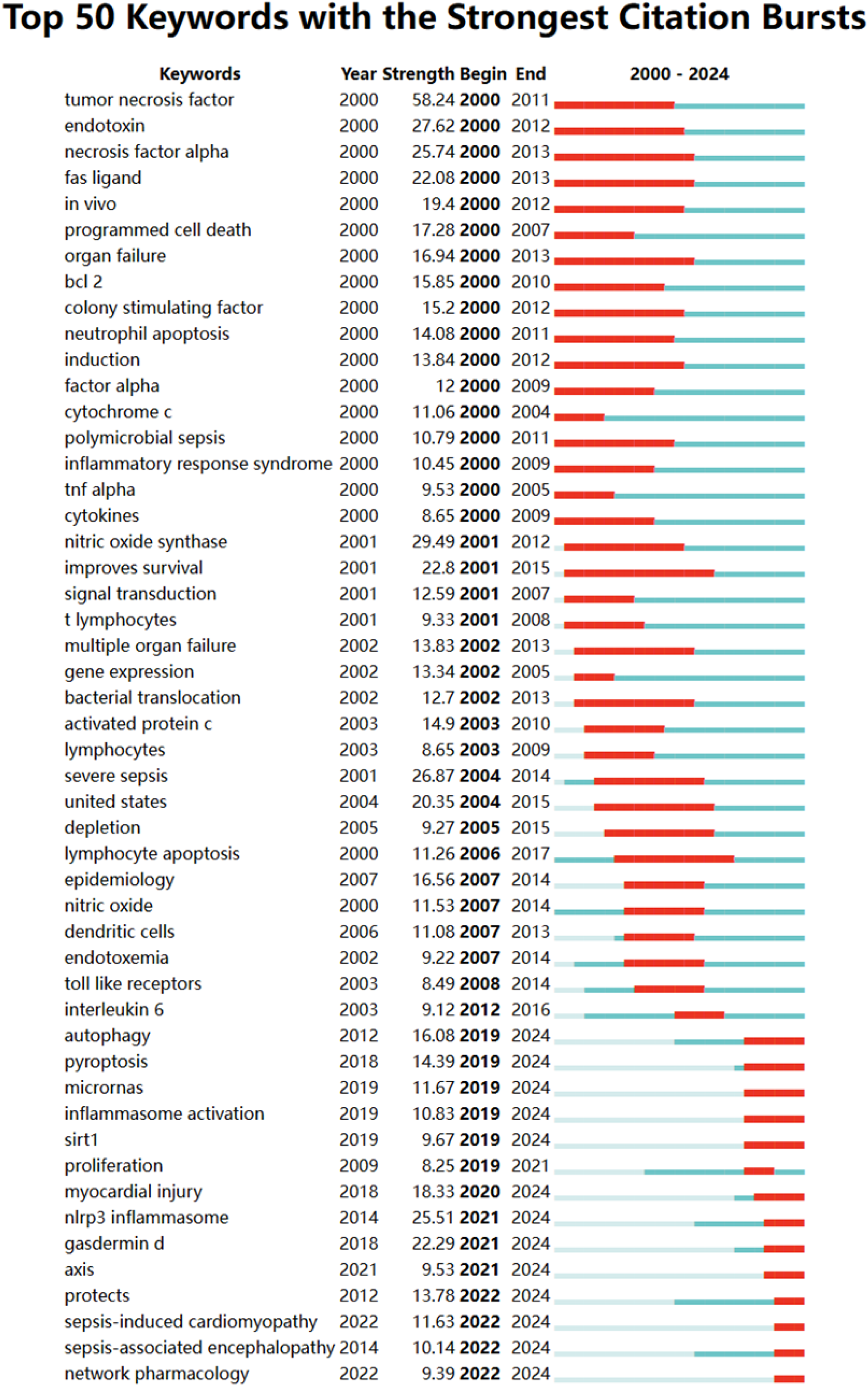
The top 50 keywords with the greatest burst intensity

#### 3.2.3. Reference burst

Through meticulous calculation, we have identified 629 groundbreaking articles. Table 3 presents the top 30 most frequently cited references spanning the period from 2000 to 2024. Singer M (2016) was the article with the first citation burst and it triggered a lot of interest when it published^22^. The Society of Critical Care Medicine and the European Society of Intensive Care Medicine convened a working group (n = 19) with expertise in sepsis pathobiology, clinical trials, and epidemiology to evaluate the definitions of sepsis and septic shock and update them as needed. These updated definitions and clinical criteria should replace previous definitions, provide greater consistency for epidemiological studies and clinical trials, and facilitate earlier identification and more timely treatment of patients with sepsis or those at risk of developing sepsis. Rudd KE had a burst strength of 67.95 from 2022 to 2024^23^. The study, which pooled data on the global burden caused by sepsis, showed a 37.0% decrease in age-standardized sepsis incidence and a 52.8% decrease in mortality from 1990 to 2017. However, sepsis incidence and mortality vary widely across regions, with the highest burden in sub-Saharan Africa, Oceania, South Asia, East Asia and South-East Asia. The Pathophysiology and Treatment of Sepsis^29^ is the third most frequently cited article. Once published, it aroused wide attention and lasted for 5 years from 2003 to 2008. The article argues that sepsis is the leading cause of death in critically ill patients in the United States. However, the individual host response to sepsis is variable, depending on the patient’s immune response, age, nutritional status, comorbid conditions, as well as the virulence of the pathogen and the size of the inoculator. This review explores the evolution of the concept of sepsis and discusses new and potential therapies. Recent clinical advances include treatment with activated protein C, strict control of blood sugar, and early goal-directed therapies to treat cellular hypoxia. Future therapies may focus on modulating the immune response according to the characteristics of a particular pathogen, the genetic characteristics of the patient and the length of the disease.

**Table 3.**
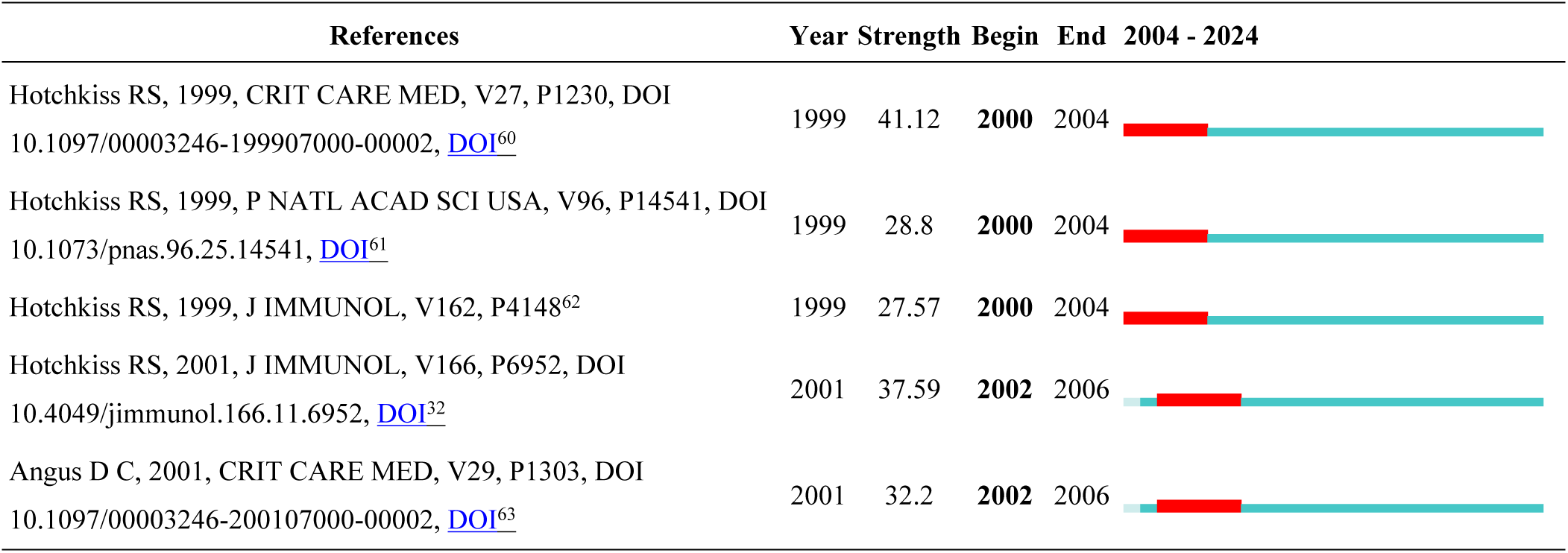
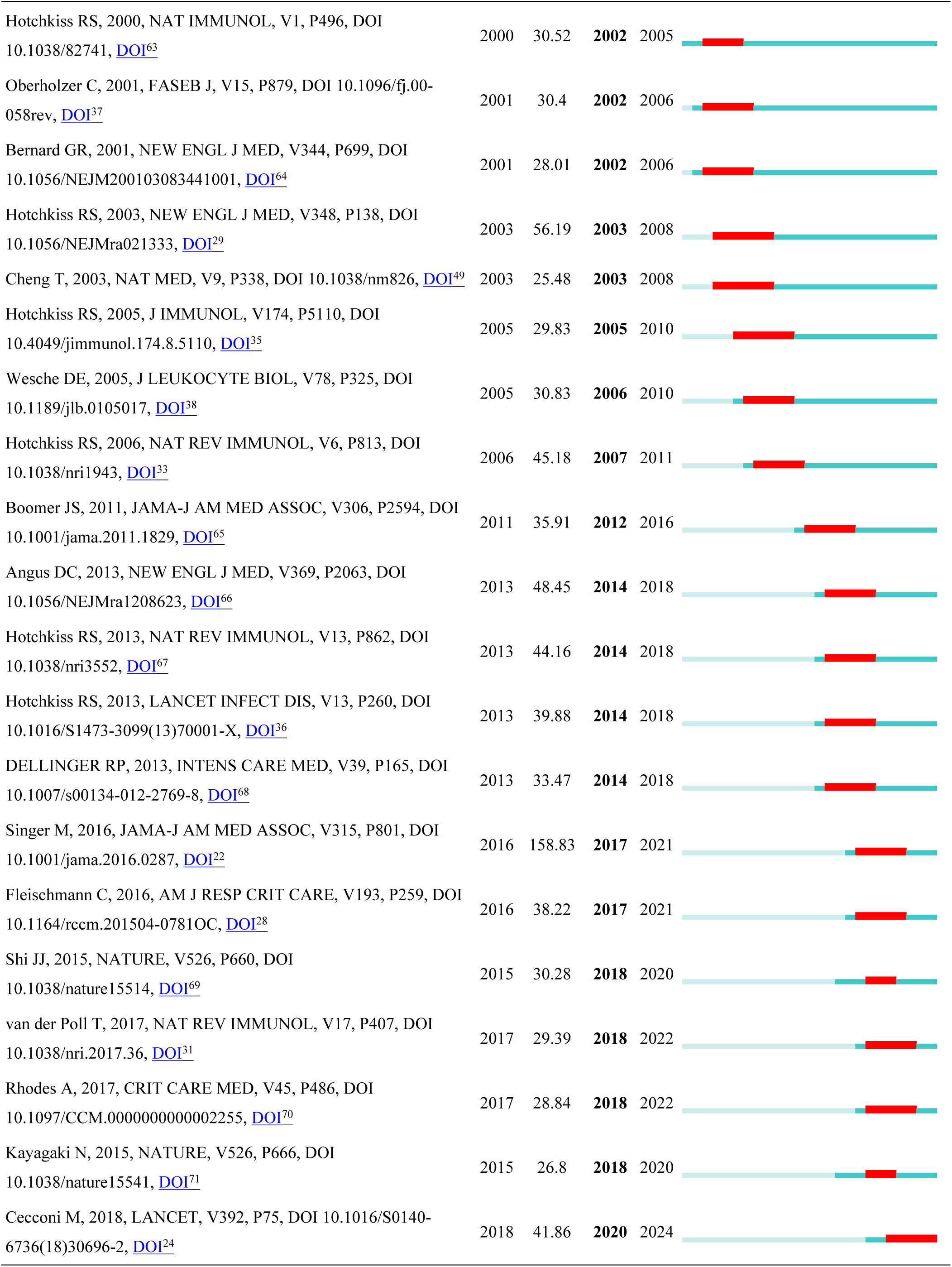
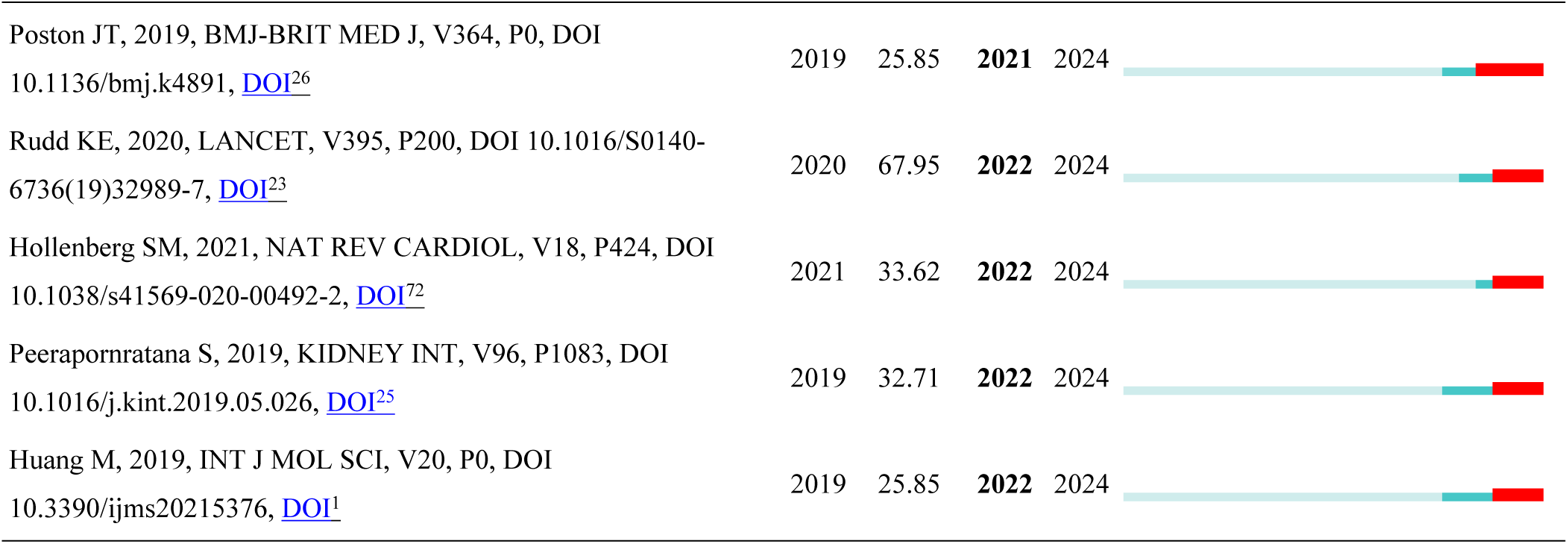
The references with citation bursts at different period.

Starting from 2024, there are 127 explosive articles, and the top 20 articles in the Power Index are shown in Table 4. Of these, 16 are “review” and 4 are “article”. All of these articles entered a citation burst period as soon as they were published that year or the following year.” The review has guiding significance for the study of cell death in sepsis, and the article has great reference value for the application of cell death in sepsis. It also reminds researchers of sepsis to pay more attention to this information about cell death.

**Table 4.**
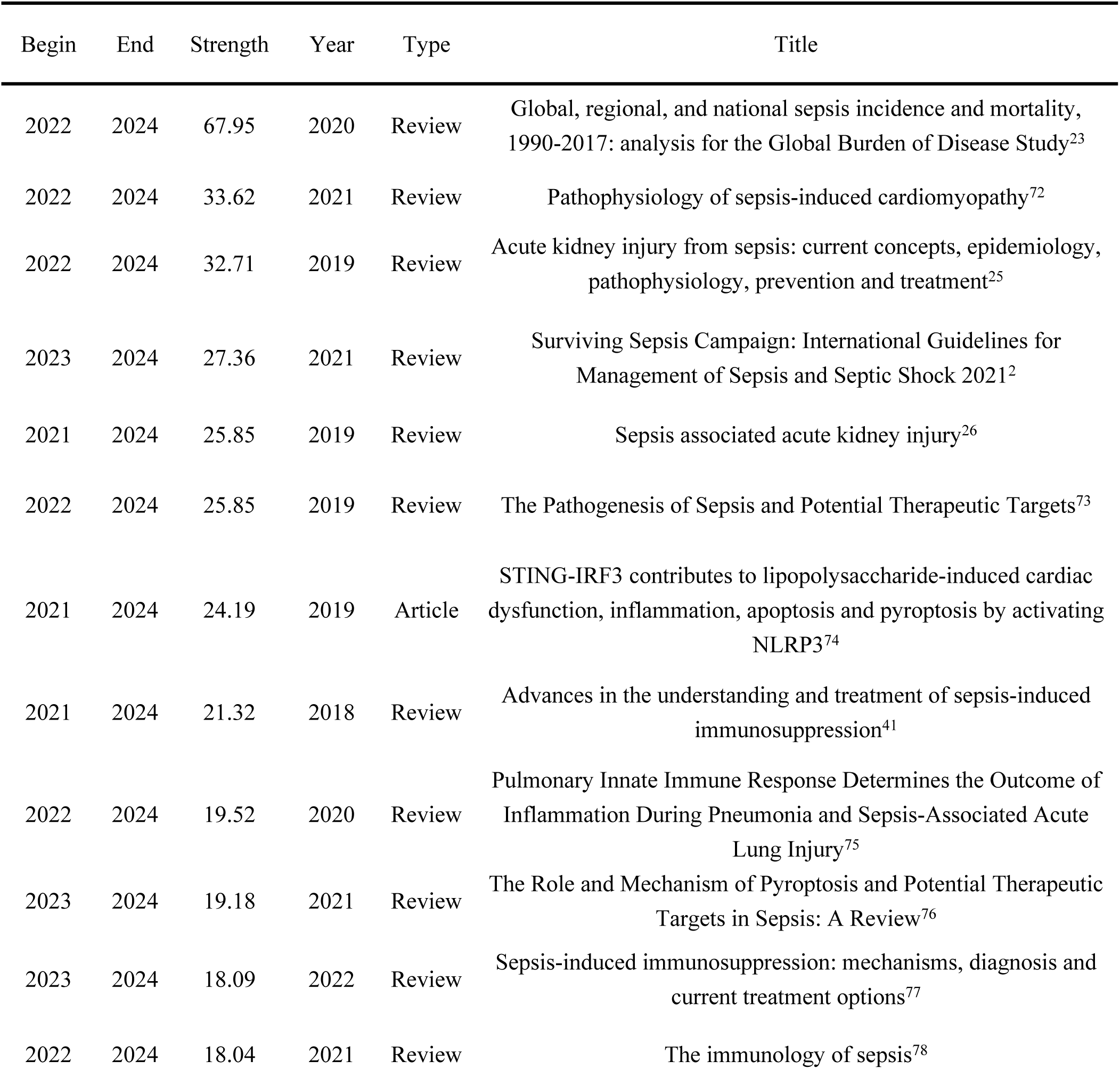
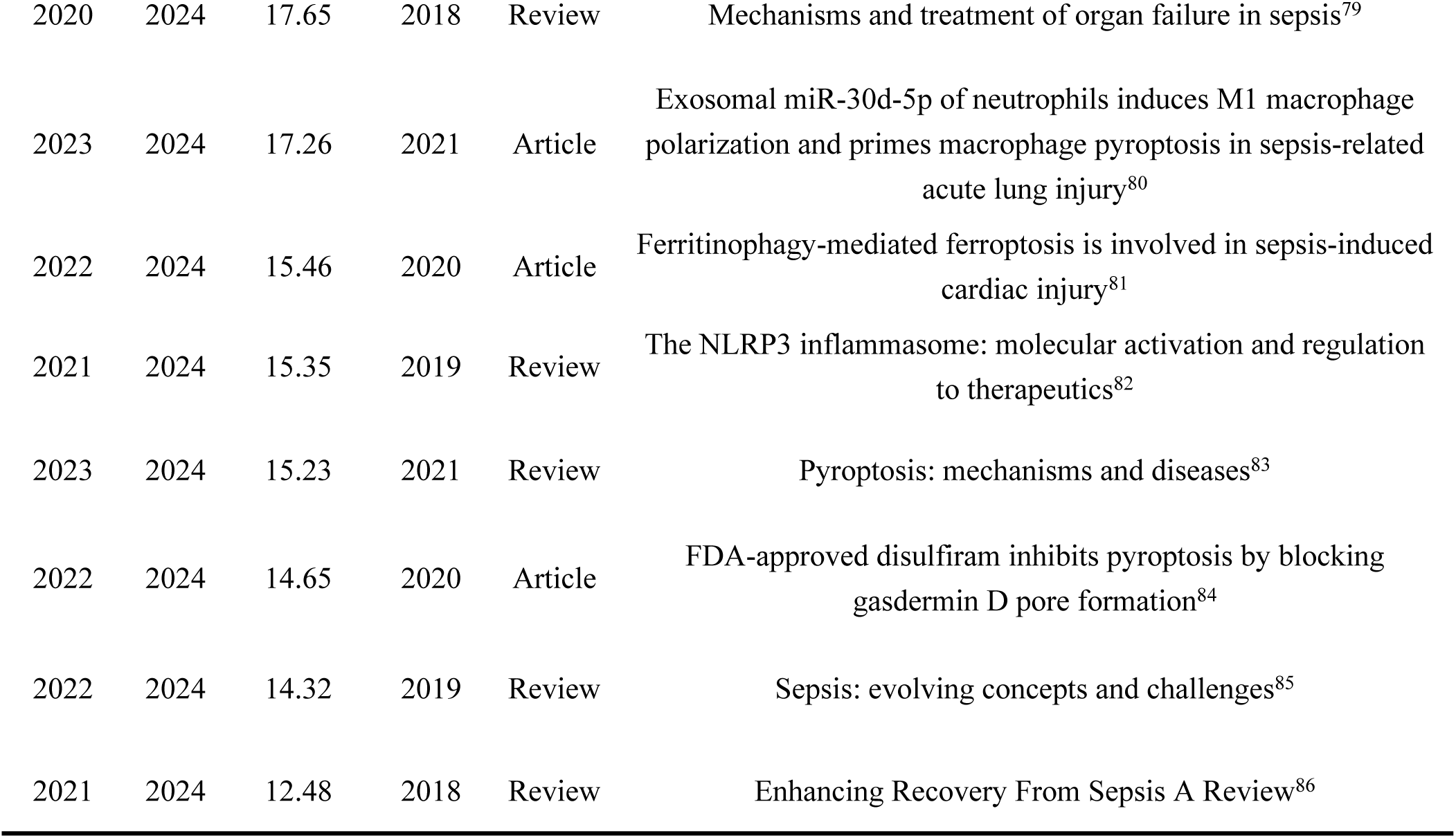
The references with citation bursts from beginning to 2024.

| Begin | End | Strength | Year | Type | Title |
| --- | --- | --- | --- | --- | --- |
| 2022 | 2024 | 67.95 | 2020 | Review | Global, regional, and national sepsis incidence and mortality, 1990-2017: analysis for the Global Burden of Disease Study<sup>23</sup> |
| 2022 | 2024 | 33.62 | 2021 | Review | Pathophysiology of sepsis-induced cardiomyopathy<sup>72</sup> |
| 2022 | 2024 | 32.71 | 2019 | Review | Acute kidney injury from sepsis: current concepts, epidemiology, pathophysiology, prevention and treatment<sup>25</sup> |
| 2023 | 2024 | 27.36 | 2021 | Review | Surviving Sepsis Campaign: International Guidelines for Management of Sepsis and Septic Shock 2021<sup>2</sup> |
| 2021 | 2024 | 25.85 | 2019 | Review | Sepsis associated acute kidney injury<sup>26</sup> |
| 2022 | 2024 | 25.85 | 2019 | Review | The Pathogenesis of Sepsis and Potential Therapeutic Targets<sup>73</sup> |
| 2021 | 2024 | 24.19 | 2019 | Article | STING-IRF3 contributes to lipopolysaccharide-induced cardiac dysfunction, inflammation, apoptosis and pyroptosis by activating NLRP3<sup>74</sup> |
| 2021 | 2024 | 21.32 | 2018 | Review | Advances in the understanding and treatment of sepsis-induced immunosuppression<sup>41</sup> |
| 2022 | 2024 | 19.52 | 2020 | Review | Pulmonary Innate Immune Response Determines the Outcome of Inflammation During Pneumonia and Sepsis-Associated Acute Lung Injury<sup>75</sup> |
| 2023 | 2024 | 19.18 | 2021 | Review | The Role and Mechanism of Pyroptosis and Potential Therapeutic Targets in Sepsis: A Review<sup>76</sup> |
| 2023 | 2024 | 18.09 | 2022 | Review | Sepsis-induced immunosuppression: mechanisms, diagnosis and current treatment options<sup>77</sup> |
| 2022 | 2024 | 18.04 | 2021 | Review | The immunology of sepsis<sup>78</sup> |
| 2020 | 2024 | 17.65 | 2018 | Review | Mechanisms and treatment of organ failure in sepsis<sup>79</sup> |
| 2023 | 2024 | 17.26 | 2021 | Article | Exosomal miR-30d-5p of neutrophils induces M1 macrophage polarization and primes macrophage pyroptosis in sepsis-related acute lung injury<sup>80</sup> |
| 2022 | 2024 | 15.46 | 2020 | Article | Ferritinophagy-mediated ferroptosis is involved in sepsis-induced cardiac injury<sup>81</sup> |
| 2021 | 2024 | 15.35 | 2019 | Review | The NLRP3 inflammasome: molecular activation and regulation to therapeutics<sup>82</sup> |
| 2023 | 2024 | 15.23 | 2021 | Review | Pyroptosis: mechanisms and diseases<sup>83</sup> |
| 2022 | 2024 | 14.65 | 2020 | Article | FDA-approved disulfiram inhibits pyroptosis by blocking gasdermin D pore formation<sup>84</sup> |
| 2022 | 2024 | 14.32 | 2019 | Review | Sepsis: evolving concepts and challenges<sup>85</sup> |
| 2021 | 2024 | 12.48 | 2018 | Review | Enhancing Recovery From Sepsis A Review<sup>86</sup> |

### 3.3. Emerging trends and new developments

#### 3.3.1. The temporal variation of keyword clusters

The keywords exhibit close interrelations, allowing them to form distinct clusters based on their correlations. Recognizing these clusters provides a more intuitive understanding of the popular subfields within sepsis research related to cell death. We divided the past two decades into four stages, each spanning six years. Figure 6 displays the snapshots of keyword clustering for each stage. In the first cluster (2000-2006), 578 articles were analyzed, yielding seven clusters, including #0 inhibition, #1 improves survival, and #2 septic shock (Figure 6A). The second cluster (2007- 2012) encompassed 963 articles, resulting in eight clusters, such as #0 inhibition, #1 activated protein C, and #2 improves survival (Figure 6B). The third cluster (2013-2018) analyzed 1483 articles, producing eight clusters, including #0 cell death, #1 oxidative stress, and #2 immunosuppression (Figure 6C). The fourth cluster (2019-2024) included 3141 articles, generating eight clusters: #0 oxidative stress, #1 pyroptosis, #2 sepsis-associated encephalopathy, among others (Figure 6D). Compared to the previous 15 years, classical topics such as septic shock and induced apoptosis remain prevalent. Additionally, emerging research clusters such as #0 oxidative stress, #1 pyroptosis, #2 sepsis-associated encephalopathy, #3 acute kidney injury, #4 immunosuppression, #5 necroptosis, #7 lung injury, and #8 extracellular vesicles have garnered increasing attention from researchers.

**Figure 6.**
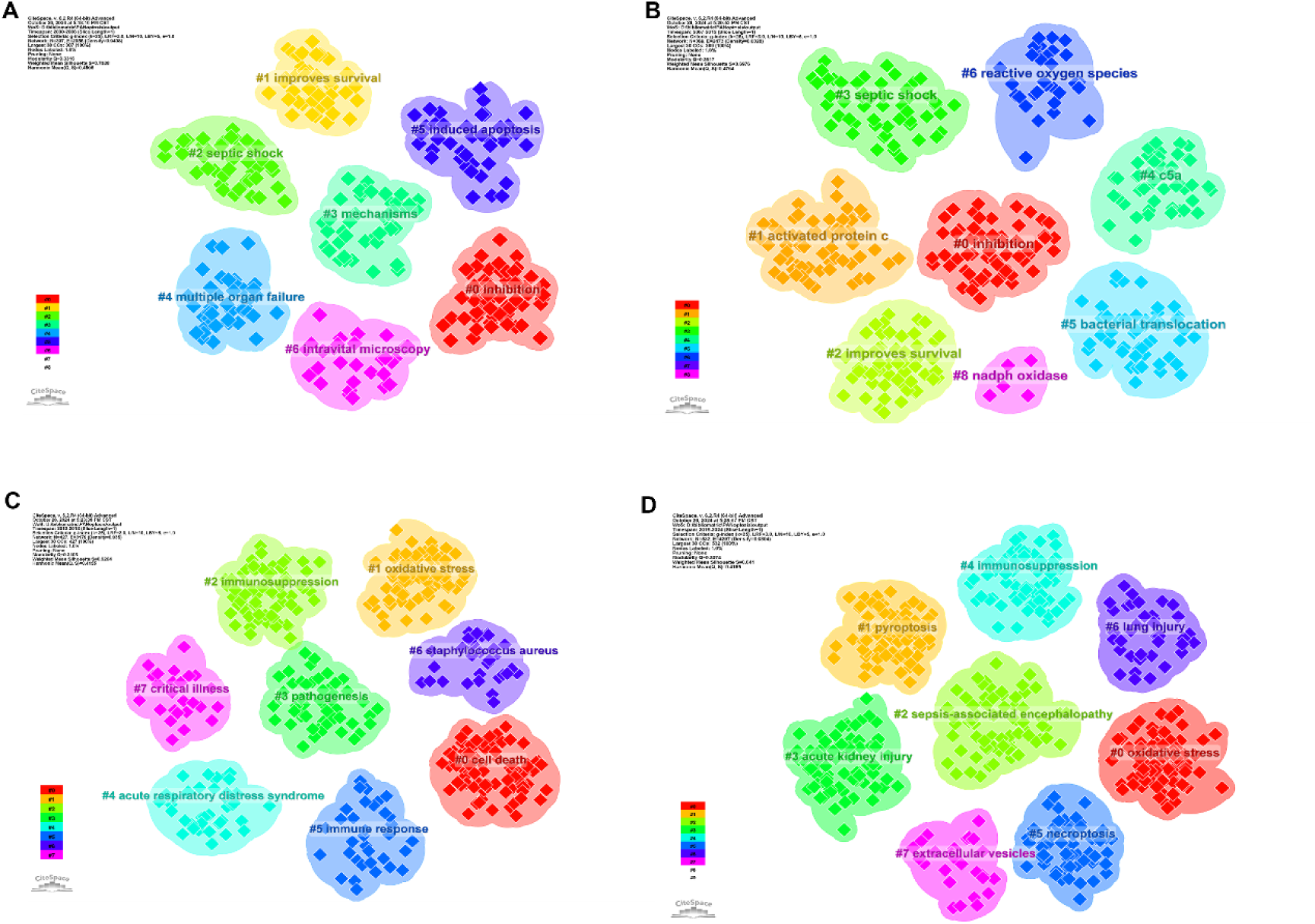
The keyword clusters snapshots in four periods

Our analysis of the literature pertaining to the emerging clusters reveals that these clusters are aimed at exploring novel techniques and methodologies. Specifically, the cluster labeled 0# oxidative stress encompasses 92 articles focused on oxidative stress in sepsis. Clusters 1# pyroptosis and 5# necroptosis, with 87 and 65 articles respectively, detail studies on various forms of cell death in sepsis. Cluster 2# sepsis-associated encephalopathy includes 76 articles focused on sepsis-related complications. Clusters 3# acute kidney injury and 7# lung injury comprise 75 and 38 articles, respectively, addressing injuries associated with sepsis. Cluster 4# immunosuppression collects 71 articles related to immunosuppression. Cluster 8# extracellular vesicles comprises 26 articles examining the role of extracellular vesicles in sepsis. Table S3 (Supplementary Materials) presents detailed data for the fourth cluster (2018-2024). The “representative keywords within the cluster” facilitate the identification of the most recent stage (2018-2024) of cell death research within the core area of sepsis.

#### 3.3.2. The keyword alluvial flow visualization

As illustrated in Figure 7, the associated keywords can be integrated into distinct research modules. Through the reconfiguration of these keywords, the research modules may differentiate or converge over various time periods, ultimately leading to the formation of novel research modules.

**Figure 7.**
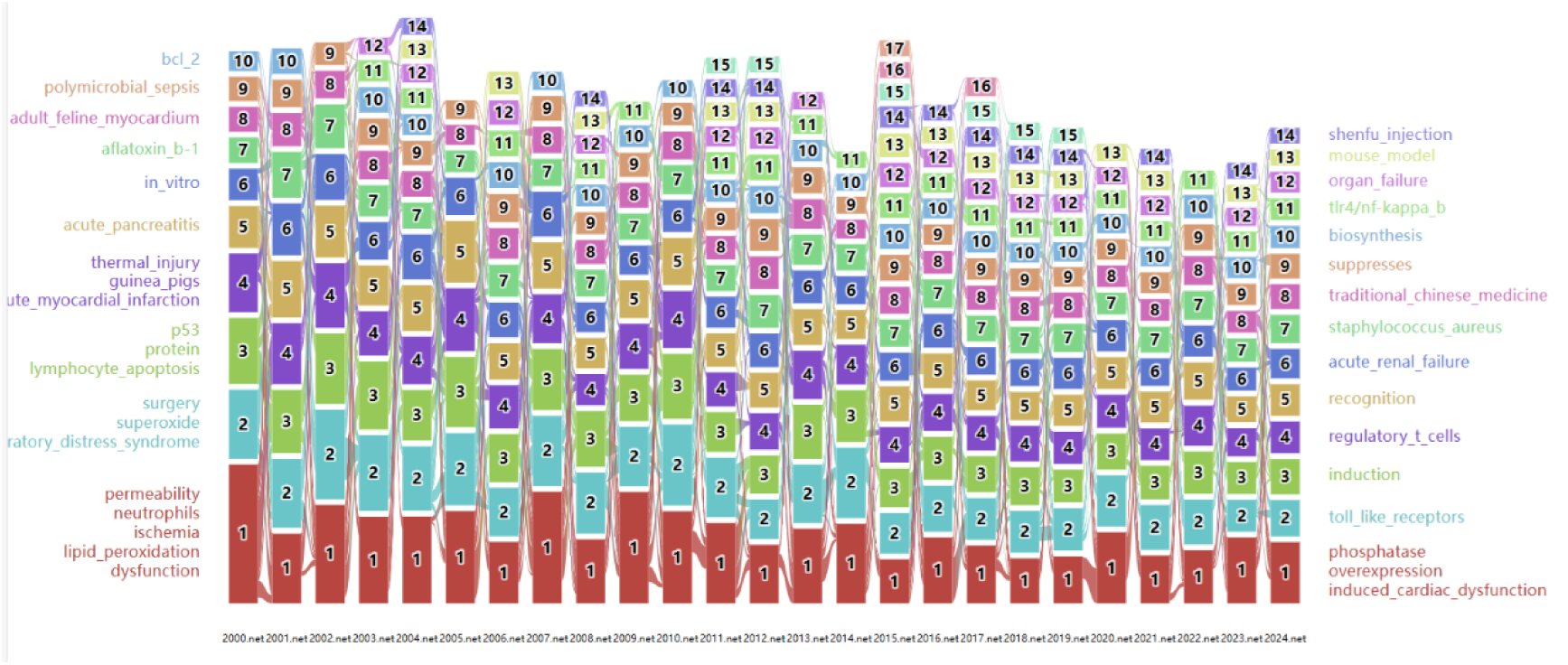
The keyword alluvial flow visualization

Over the past quarter-century, certain traffic keywords have exhibited robust vitality, some evolving into new research trends while others fading into the annals of the research domain. Table S4 (Supplementary Materials) enumerates the keywords associated with the top five modules exhibiting the highest annual traffic. Notably, the keywords constituting Module 1 in 2024, which have either diverged or converged within this research arena, constitute the most prominent research tributaries (highlighted in red). This underscores Module 1 as the most enduring research module. Furthermore, we have delineated all keywords pertaining to the top six modules in 2024 in Figure 8. Module 1, designated as “phosphatase,” encompasses 11 keywords including overexpression, induced_cardiac_dysfunction, and heart (Figure 8A). Module 2, termed “toll_like_receptors,” comprises four keywords such as mitophagy, barrier, and models (Figure 8B). Module 3, named “induction,” encompasses 11 keywords including ferroptosis, migration, cardiomyocyte_apoptosis, and differentiation (Figure 8C). Module 4, designated as “regulatory_t_cells,” consists of five keywords including immune system and risk (Figure 8D). Module 5, termed “recognition,” includes 11 keywords such as lung, cleavage, and neuroinflammation (Figure 8E). Module 6, designated as “acute_renal_failure,” comprises nine keywords including ginsenoside_rg1, growth, cell_apoptosis, and homeostasis (Figure 8F). These modules may represent emerging trends in the field of sepsis-related cell death over the next five years or beyond.

**Figure 8.**
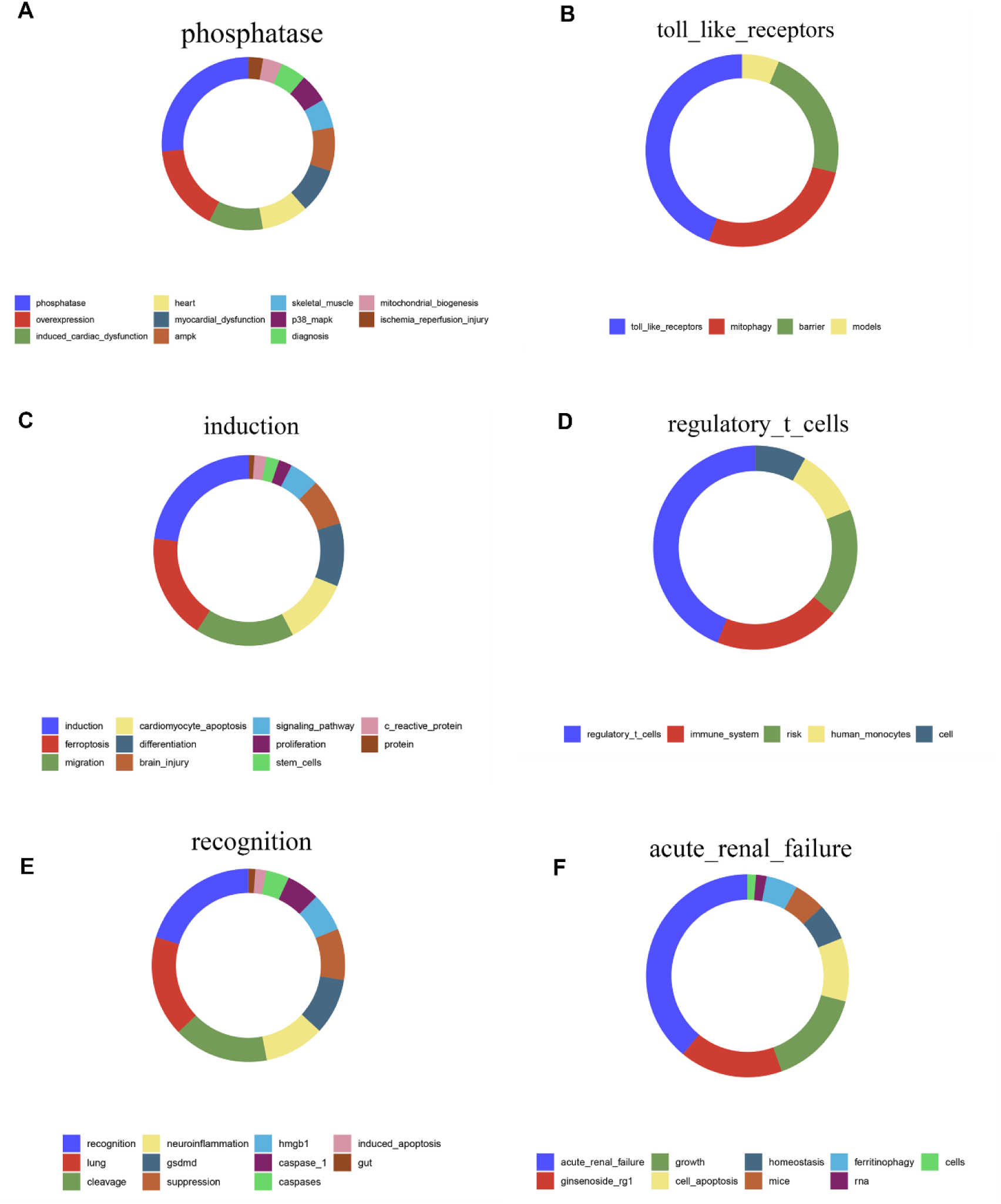
The keywords of the top 6 modules in 2024.

#### 3.3.3. The timeline visualization of references

A timeline visualization based on the time span of the citation can predict which topics are emerging, which are classic, and which are relatively outdated. The timeline diagram of cell death in sepsis studies consists of 16 groups arranged from top to bottom by size at a given time (Figure 9A). Among them, # 5 chemokines, # 6 lymphocyte, # 7 hippocampus, # 8 activated protein C, # 9 renal insufficiency, # 10 foxo3a, # 14 zymomonas mobilis, #15 endothelin-converting enzyme were classic topics, which may not be the latest topic but inextricably linked to other clusters. # 0 caspase, #11 neuromediators, and #12 adrenal were relatively outdated topics that have little connections with other clusters and no follow-through in their own timeline. #1 immunosuppression, #2 sepsis-induced cardiomyopathy, #3 pyroptosis, #4 acute kidney injury and #13 covid-19 were emerging topics. Because they have been active in the timeline since their emergence to the present, this indicates that these areas will become future research hotspots. Table S5 (supplementary material) shows the emerging clusters in more details. In addition, some classic papers (large nodes with red circles) have played a very important role in advancing the subfield (Figure 9B). An article published by van der Poll T in 2017^31^, belonging to cluster #1 with the co-citation frequency of 96. The article shows that in sepsis, the immune response triggered by the invading pathogen cannot be returned to a state of equilibrium, which ultimately leads to a pathological syndrome characterized by persistent excessive inflammation and immunosuppression. The key to the clinical development of new therapies for sepsis is the selection of patients based on biomarkers and/or functional deficits that provide specific information about the expression or activity of therapeutic targets. Li N (2019, Redox Biol)^74^, belongings to cluster #2 with the co-occurrence frequency of 114, evaluated that STING can bind to type I interferon (IFN) regulatory factor 3 (IRF3) and phosphorylate IRF3. Phosphorylated (P-) IRF3 is subsequently translocated into the nucleus and increases the expression of NOD-like receptor protein 3 (NLRP3). Isolated TXNIP can directly interact with NLRP3 in cytoplasm and form inflammasome, which eventually causes myocardial cell injury. Deng M (2018, Immunity)^50^, belongings to cluster #3, with the co-occurrence frequency of 69, demonstrated that HMGB1 released by hepatocytes binds to LPS and targets it into lysosomes of macrophages and endothelial cells via the Advanced glycation end product Receptor (RAGE). Subsequently, HMGB1 permeates the phospholipid bilayer in the acidic environment of the lysosome. This causes LPS to leak into the cell membrane and activate caspase-11. Peerapornratana S^25^ (2019, Kidney Int), belongings to cluster #4, which demonstrated that a novel biomarker of renal stress and injury was validated for risk prediction and early diagnosis of acute kidney injury in the context of sepsis. Microvascular dysfunction, inflammation, and metabolic reprogramming are three fundamental mechanisms that may play a role in the development of S-AKI. Fan, E. K. Y ^87^ (2018, Respir Res), belongings to the cluster #13, which demonstrated that the death of alveolar macrophages (AM) plays an important role in the development of lung inflammation by affecting other populations of immune cells in the lung. Cell death and tissue inflammation form a positive feedback loop, ultimately leading to increased inflammation and disease development. Pharmacological regulation of AM death signal can be used as a rational treatment strategy for ALI/ARDS. We further statistical the citation distribution of these five articles in recent years (Figure 9C), and can predict that these articles may be mentioned again in the next few years.

**Figure 9.**
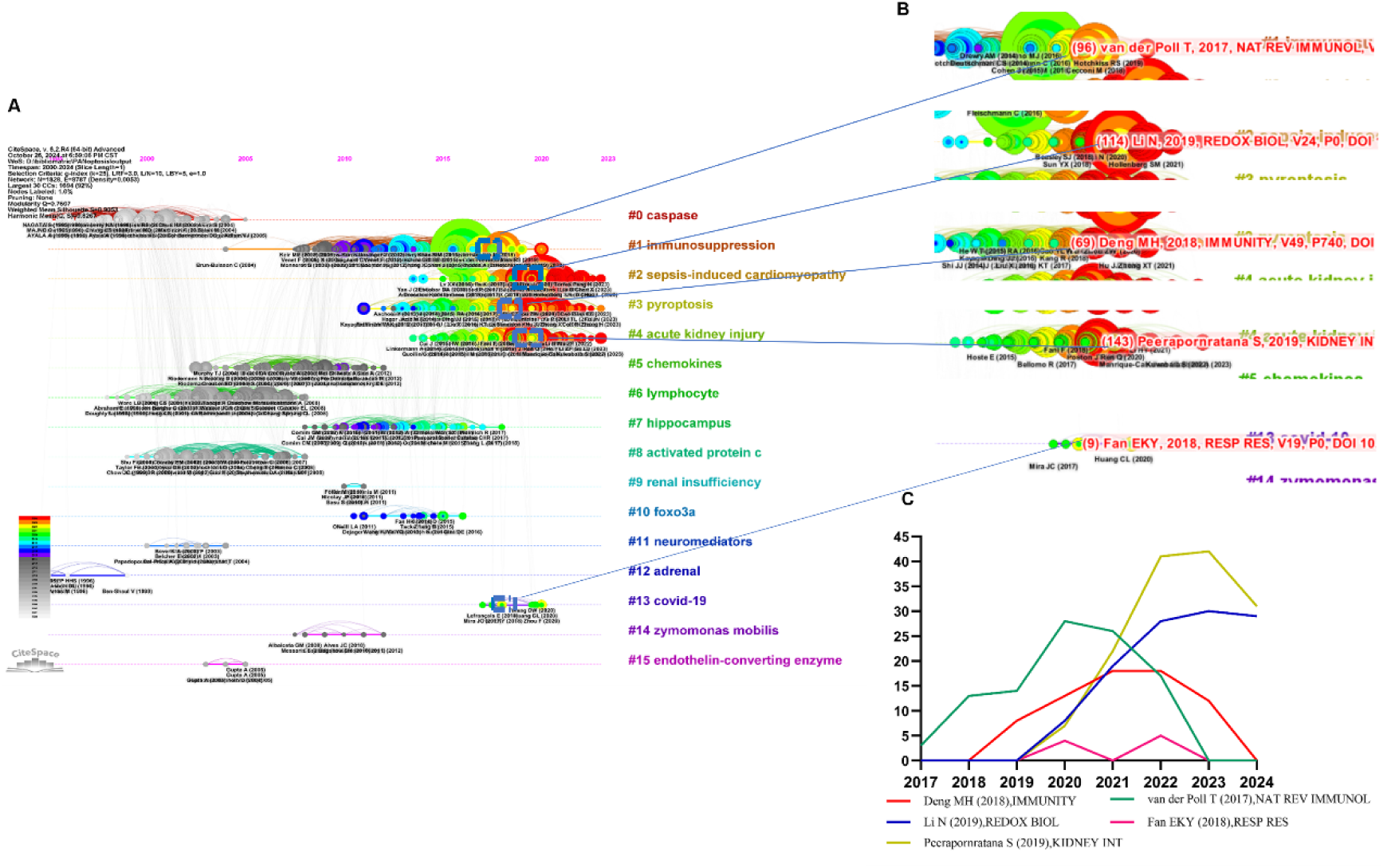
The reference clusters map. A: the citation timeline visualization. B: The burst citation in #1, #2, #3, #4, #13. C: citation frequency distribution of the burst citation.

## 4. Discussion

In our research, we employed some scientific tools to conduct a comprehensive review of the structural and temporal characteristics of publications within the field of PANoptosis in sepsis spanning from 2000 to 2024. To the best of our knowledge, this is the first bibliometric review, summary and outlook of this field. It is unequivocally evident that the field of PANoptosis in sepsis remains in a vibrant and active stage, characterized by a substantial surge in publication volume, extensive and intensive scientific collaborations, as well as a tightly interconnected citation network.

### 4.1 General information

The application of bibliometric analysis to assess trends and advances in specific fields has gained widespread popularity. However, although it is a hot area of current research, no studies have used bibliometrics to analyze the trends and hot spots of PANoptosis in sepsis research. Based on the scientific atlas of bibliometrics, this paper reviews the structure and temporal characteristics of publications in the field of sepsis panapoptosis from 2000 to 2024. Our study found that research on panapoptosis in sepsis is expanding rapidly, especially after 2018, with a further increase in the number of publications reaching a peak in 2022. It is worth noting that the number of publications of shock journals, which are generally considered to be the most representative journals, has reached 256. Other major journals, such as International Immunopharmacology, Frontiers in Immunology, Critical Care Medicine, are also involved in the publication of sepsis panapoptosis. Therefore, we believe that more cutting-edge research on relevant topics can be obtained from the above excellent journals.

After surveying the current state of research cooperation between countries around the world, our findings show that the intensity of cooperation between China and the United States, as well as other countries, is very high. This partly explains why China and the United States can publish more articles in this field. At the same time, we further shift the research perspective from the country to the specific research institution. Among these institutions, Central South University has been the most active in publishing articles in this field, demonstrating significant investment in scientific research and remarkable achievements. At the same time, the results in the authors’ collaboration map indicate significant collaboration among research institutions, indicating that teamwork is an effective academic communication strategy^88^ and that scholars and institutions around the world actively contribute their expertise to removing academic barriers and promoting academic collaboration and exchange. We firmly believe that outstanding scholars can contribute more to the hot research in this field, which will play a key role in further exploring the pathogenesis and therapeutic targets of sepsis.

While hot topics in the field change over time, the latest keywords with citation burst results show that tumor necrosis factor, nitric oxide synthase, endotoxin is the three key words with high outbreak intensity, among which nlrp3 inflammasome, gasdermin d, myocardial injury and autophagy still have high outbreak intensity in 2024, which may become future research hotspots. The latest citation explosion literature is also studied around these hotspots. These results are also reflected in the keyword cluster map and the reference timeline map. In addition, the results of keyword timeline visualization show that chemokines, lymphocyte, hippocampus and other keywords are classic themes, while immunosuppression, sepsis-induced cardiomyopathy, pyroptosis, acute kidney injury. covid-19 is an emerging topic.

### 4.2 Hotspots and Frontiers

#### 4.2.1 The Crosstalk Between Pyroptosis, Apoptosis and Necroptosis

An increasing number of studies have shown that the three pathways involved in PANoptosis, namely apoptosis, pyrodeath and necrotic apoptosis, can be activated together in the same cell. Moreover, it has been shown that although the three pathways operate in parallel, they can be cross-regulated with each other by a multi-protein complex called PANoptosome, which promotes the death of inflammatory cells. In a rat model of septic-associated encephalopathy, apoptosis inhibitors inhibit pyrodeath and activate necrotic apoptosis. Pyroptosis inhibitors can inhibit pyroptosis and apoptosis, but can activate necrotic apoptosis^89^. However, when necrotic apoptosis is inhibited, both pyrodeath and apoptosis can be activated^89^. Among them, caspase-8, as one of the earliest discovered Bridges between different cell death types, is an important regulator of cell pyroptosis, apoptosis and necroptosis^90^. CASP8 is not only the initial caspase for exogenous apoptosis, but also inhibits RIPK3 and MLKL-mediated necroptosis^91,92^. Meanwhile, other studies have shown that caspase-8 is a key component required for transcriptional initiation and activation of both typical and atypical NLRP3 inflammasome in mice^91,93^. When TAK1 is inhibited, its activation leads to the cleavage of GSDMD and GSDME in mouse macrophages^90,94^. In addition to playing a key role in pyrodeath, CASP1 can activate CASP7, promote its translocation to the nucleus, and cut PARP1 on the promoter of a subunit of the NFKB1 target gene. If downstream of CASP1, such as GSDMD, is missing, apoptotic pathways can be induced by splitting CASP3^95–97^. Necroptosis usually occurs in the absence of CASP8, which can cut RIPK1 or RIPK3^98,99^. When RIPK1 is expressed in low or insufficient amounts, cells die due to apoptosis^100^. So far, CASP8 plays an important transforming role as the intersection of pyrodeath, apoptosis, and necroptosis^9^. In addition, as the study progressed, the researchers confirmed that the activity of the NLRP3 inflammasome is a driver of inflammation in MLKL-dependent diseases. Under certain conditions, the necrotic apoptotic signal can activate the RIPK3-MLKL-Caspase-1 signal axis, leading to the maturation and release of IL-1β, and this process is independent of GSDMD^101–103^.

#### 4.2.2 Role of PANoptosis in the pathogenesis of sepsis

Although pyroptosis, apoptosis, and necroptosis have all been studied in the pathogenesis of sepsis, their interactions at the molecular level have been largely unknown. Apoptosis-mediated cell death often occurs in heart, kidney, and other organ failure during sepsis. Pyroptosis is usually associated with fatal sepsis^74,104,105^. Necroptosis has also been observed in renal damage caused by mitochondrial dysfunction in sepsis^106^. PANoptosis, a mode of programmed cell death widely reported in oncologic diseases^107^, has been less studied at the molecular level in sepsis. The study found that bacteria can induce the development of PANoptosis by ZBP1, interferon regulatory factor 1, or other risk genes^7,108,109^. The current study shows that PANoptosis is an important mechanism for the body to fight infection. Sepsis triggered by infection triggers a systemic inflammatory response and often leads to widespread cell death. Studies have shown that PANoptosis is simultaneously activated in rats with sepsis related encephalopathy^89^, and inhibiting the occurrence of necrotic apoptosis can normalize the apoptosis of nerve cells, thereby protecting neuronal damage^110^. Recent studies have shown that inhibiting STING pathway mediated panapoptosis can improve the occurrence of lung injury in sepsis^13^. As a new cell death mode, PANoptosis integrates key molecular mechanisms of apoptosis, necroptosis and pyroptosis to form a unique regulatory network. Key node molecules, such as ZBP1, RIPK1, RIPK3 and caspase-8, cooperate or restrict each other to accurately regulate cell fate. In the sepsis environment, pathogen infection and inflammatory mediators stimulate intracellular stress signals to converge to the regulatory hub of panapoptosis. Depending on cell type, stimulation intensity and microenvironment differences, cells are determined to go to panapoptosis or other death paths, thus affecting the local inflammatory microenvironment, immune cell function and tissue repair and regeneration potential. A key piece in the complex pathophysiological puzzle of sepsis. Therefore, understanding the interaction between different cell death modes in sepsis is of great significance for further understanding their crosstalk mechanism and pathological process in sepsis.

## 5. Limitations

Although the current research of PANoptosis in the field of sepsis has achieved many stage results, many difficulties remains. On the one hand, the fine regulation details of the PANoptosome signaling pathway are still unclear, and the interaction mechanism between many key molecules has not been fully clarified. For example, the precise activation mode of different components of the PANoptosome complex under different cell types and stimulation conditions needs to be further explored. There are a large number of unknown nodes in the synergistic or antagonistic regulatory network among various catalytic effectors, which greatly limits the deep understanding of the core mechanism of PANoptosis and hinders the accurate design of targeted intervention strategies. On the other hand, the existing cell death detection technology still has shortcomings in accurately distinguishing PANoptosis from other cell death modes. Traditional morphological observation and biochemical index detection are difficult to meet the specific recognition needs of PANoptosis in complex vivo environment, and lack of highly sensitive and highly specific molecular markers, which is easy to cause misjudgment of experimental results. Evaluation of the true occurrence and effect of interference on PANoptosis. In addition, given that PANoptosis involves complex signal networks and molecular regulation, the development of efficient, low-toxicity and highly specific targeted therapeutic drugs is a challenge. When drug molecules interfere with the process of PANoptosis, it is easy to cause the imbalance of other physiological functions such as immunity and metabolism. So it is an urgent problem to overcome about how to optimize drug design to achieve accurate regulation of PANoptosis while maintaining body homeostasis Become an urgent problem to overcome.

## 6. Conclusions

Despite the challenges, PANoptosis research in the field of sepsis is still emerging and promising for the future. On the one hand, with the rapid development of multi-omics technology, the integration of genomics, proteomics, metabolomics and single-cell sequencing technology is expected to comprehensively and accurately analyze the dynamic molecular map of PANoptosis in different stages of sepsis and different cell types, accurately identify key regulatory nodes and molecular markers, and provide a “navigation map” for targeted therapy. On the other hand, based on the in-depth understanding of the molecular mechanism of PANoptosome, it is possible to develop new drugs that specifically target key proteins of the PANoptosome complex or its upstream and downstream signaling molecules. These drugs can not only accurately interfere with the process of PANoptosome, but also minimize the interference with normal physiological functions of the body. To bring more effective treatment options to patients with sepsis. In addition, the combination of multiple therapeutic strategies, such as the synergistic application of targeted PANoptosis therapy with immunomodulatory therapy, blood purification technology, intestinal flora intervention, etc., is expected to reshape the immune balance, metabolic homeostasis and microecological environment of the body in sepsis from multiple angles and levels, break the current deadlock in sepsis treatment, and significantly improve the survival rate and quality of life of patients.

## Ethics approval and consent to participate

Not applicable

## Availability of data and materials

The authors commit to making the raw data underlying the conclusions of this manuscript freely accessible, without unnecessary restriction, to any researcher who meets the requisite qualifications.

## Authors’ contributions

ZHL: Conceptualization, Data curation, Methodology, Validation, Writing-original draft. DN: Software, Data acquisition. LLY: Data curation, Methodology, Formal analysis. QDQ: Data curation, Methodology, Validation. CBY: Formal analysis, Investigation, Writing-review & edit. RL: Methodology, Formal analysis. XMG: Resources, Investigation, Software. YW: Data curation, Visualization. XYY: Investigation, Funding acquisition, Project administration, Supervision.

## Acknowledgments

Thanks to the members of our laboratory for their valuable contributions to this work.

